# Structural basis of botulinum neurotoxin serotype A1 binding to human SV2A or SV2C receptors

**DOI:** 10.1101/2022.07.06.498993

**Authors:** Fodil Azzaz, Didier Hilaire, Jacques Fantini

## Abstract

Botulinum neurotoxin A1 (BoNT/A1) is the most potent serotype in humans with the highest clinical duration. BoNT/A1 interacts with synaptic vesicle glycoprotein 2 (SV2) and gangliosides to be taken up by neurons. In this study, we present three molecular dynamics simulations in which BoNT/A1 is in complex with singly or doubly glycosylated SV2C or singly glycosylated SV2A, in a ganglioside rich (lipid raft) context. Our computational data suggest that the N-glycan at position 480 (N480g) in the luminal domain of SV2C (LD-SV2C) indirectly enhanced the contacts of the neurotoxin surface with the second N-glycan at position 559 (N559g) by acting as a shield to prevent N559g to interact with residues of LD-SV2C. The N-glycosylation at the position N573 (N573g) in the luminal domain of SV2A has a slightly lower affinity for the surface of BoNT/A1 compared to 559g because of possible intermolecular contacts between N573g and residues of the luminal domain of SV2A (LD-SV2A). In addition to the ganglioside binding site (GBS) conserved across serotypes B, E, F and G, the lipid-raft associated GT1b interacted with a structure we coined the ganglioside binding loop (GBL) which is homologous to the lipid binding loop (LBL) in serotypes B, C, D, D/C and G. Finally, we proposed a global model in which BoNT/A1 interacts with its glycosylated protein receptor, one molecule of GT1b interacting in the GBS and five molecules of GT1b interacting with the GBL and residue Y1133. These data solved the puzzle generated by mutational studies that could be only partially understood with crystallographic data that lack both a biologically relevant membrane environment and a full glycosylation of SV2.

**Brief statement:** We propose a full molecular description of the initial binding of a microbial toxin (Botulinum neurotoxin A1) to the surface of neural cells. Our model includes a protein receptor (SV2) in its native environment, i.e. the periphery of a cluster of gangliosides belonging to a membrane microdomain (lipid raft). A major outcome of our study is the elucidation of the role of the full length glycans (previously resolved by MS spectroscopy) covalently attached to the protein receptor. These data solved the puzzle generated by mutational studies that could be only partially understood with crystallographic data that lack both a biologically relevant membrane environment and a full glycosylation of SV2.

## Introduction

Clostridia strains are gram-positive, anaerobic, spore-forming bacilli that live in dust, soil, vegetation, water, and the gastrointestinal tract of animals *^1–4^*. They produce toxins which belong to the Clostridial neurotoxin family comprising tetanus toxin and the eight major serotype of botulinum neurotoxins (BoNTs) classified from A to X *^5–7^*. Among them, the serotypes A, B, E and F were reported to be associated with human botulism while the serotypes C and D are associated with animal botulism *^8, 9^*. They interact with specific membrane receptors at the neuromuscular junction of alpha-motoneurons and cause botulism by inhibiting neurotransmitter release via the cleavage of SNARE proteins comprising syntaxin 1, VAMP1-3 or SNAP-25 *^10^*. As a result, the infected subject undergoes a flaccid paralysis which can lead to death by causing respiratory failure. Of the four serotypes that infect humans, botulinum neurotoxin A (BoNT/A1) is the most potent followed by serotypes B (BoNT/B), E (BoNT/E) and F (BoNT/F) *^11^*. Notwithstanding the lethality of this agent, BoNTs are widely exploited in the medical domain to treat a broad range of neurological disorders as well as in the field of cosmetics *^12–14^*

BoNTs are composed of two domains comprising a 50 kDa light chain (LC) and a 100 kDa heavy chain (HC). The LC is the catalytic domain of BoNTs. It is associated with the HC via a disulphide bond and a set of non-covalent interactions *^15^*. The HC is divided into two subdomains, the 50 kDa N-terminal subdomain of translocation (HN) and the 50 kDa C-terminal subdomain (Hc). Hc is further divided into HCN and HCC of 25 kDa each *^16^*. All these domains and subdomains play a distinct role in the multistep mechanism of neuron intoxication. First, the Hc domain recognizes and binds its specific membrane receptors in the synaptic endings of cholinergic neurons which drive the internalization of the neurotoxin. Then, upon the acidification of the environment, the subdomain HN translocates the LC into the cytoplasmic domain to cleave its targets responsible for the neurotransmitter vesicles fusion *^17^*.

BoNT/A1 uses as protein receptors the luminal domain of the three isoforms of synaptic vesicle glycoproteins 2 (SV2s) denoted SV2A, SV2B and SV2C *^18^*. Among them, the non-glycosylated SV2C appears to be the receptor which has the highest affinity for BoNT/A1 *^18, 19^*. SV2s are predicted to consist of 12 transmembrane domains and reside in the membrane of synaptic vesicles. The luminal domain located between transmembrane domains 7 and 8 is glycosylated and displays 3 putative N-glycosylations sites for SV2A and SV2B or 5 putative N-glycosylation sites for SV2C. Three of these sites are homologous in SV2A and SV2B while two of them are unique to SV2C (at the positions N480 and N563). An experimental mapping consisting of truncating the luminal-domain of SV2C (LD-SV2C) revealed that BoNT/A1 recognizes the amino acid sequence 529 to 566 as its binding region *^18^*. The crystal structure resolved by X ray diffraction of the Hc subdomain of BoNT/A1 (Hc-BoNT/A1) co-crystalized with non-glycosylated LD-SV2C shows that BoNT/A1 binds LD-SV2C via its beta-hairpin structure and its binding is mainly mediated by backbone-backbone interactions between the beta structures *^20^*. The N-glycosylation at position N559 in the LD-SV2C enhances the interaction of BoNT/A1 to its protein receptor and to the neurotoxin surface *^21, 22^*. The disruption of the N-glycosylation site of SV2A at position N573, which is homologous to N559 in SV2C, reduced the entry of BoNT/A1, suggesting a similar mechanism of interaction between the isoforms *^23^*. The structure of Hc-BoNT/A1 co-crystalized with the LD-SV2Cg provides structural clues to the neurotoxin residues involved in the interaction with the sugars of the N-glycosylation *^22^*

On the other hand, BoNTs bind to gangliosides via a ganglioside binding site (GBS) comprising a motif conserved among the serotypes A, B, E, F, G, and tetanus neurotoxin *^24^*. Gangliosides interact with cholesterol to form condensed plasma membrane microdomains referred to as lipid rafts. These negatively charged platforms attract a wide range of cationic toxins and pathogens, and initiate their entry into target cells *^25–29^*. The addition of exogenous gangliosides increased the level of coimmunoprecipitation of BoNT/A1 with SV2s *^18^*.

Currently, no models of BoNT/A1 interacting with its membrane receptor in a ganglioside rich (lipid raft) context and respecting the global topology and the geometric constraints of a neural membrane is available. To improve our knowledge about the dynamic behind the molecular mechanism of interaction of BoNT/A1 in a lipid raft context, we performed an all-atom molecular dynamics simulation of the Hc subdomain BoNT/A1 in interaction with glycosylated membrane-embedded SV2A and SV2C in a lipid raft context.

## Material and methods

### System set-up

The full-length membrane protein SV2A and SV2C were generated via ab-initio modeling using the graphical user web-based platform Robetta (https://robetta.bakerlab.org/). The molecular docking of the Hc domain of BoNT/A1 with SV2A or SV2C was performed on the software HyperChem *^30^*. Each complex was submitted to minimization with the Polak-Ribière conjugate gradient algorithm, with the Bio+(Charmm) force field in Hyperchem and a root-mean-square (RMS) gradient of 0.01 kcal.Å-1.mol-1. The coordinates of ganglioside and cholesterol molecules composing the lipid raft were retrieved on CHARMM-GUI using the tools “glycolipid modeler” and “membrane builder” *^31, 32^*.

The glycosylation of SV2A (singly at the position N573), SV2C (singly glycosylated at position N559), and SV2C (doubly glycosylated at positions N480-N559) were generated using the tool PDB-reader available on the platform CHARMM-GUI *^33, 34^*. The lipid raft and the proteins were embedded in a membrane consisting of 50% POPC/50% cholesterol to mimic a lipid raft condition. Then, the system was solvated with TIP3P water model using the plugin “Add Solvation Box” and neutralized (with Na^+^ and Cl^-^ ions at a final concentration of 0.15 mol/L) using the tool “Add Ion” available in VMD software *^35^*.

### Molecular dynamics simulations

The systems were simulated using NAMD 2.14 for Windowsx64 *^36^* coupled with the all-atom Force Field Charmm36m *^37^*. The simulation was performed on a Dell local workstation with a mean calculation speed of 14 hours to simulate 1ns.The cutoff for the calculation of non-bonded interaction was set at 12 Å. The PME calculation algorithm was used to calculate long range electrostatic interactions. The covalent bonds involving hydrogen atoms were constrained using the algorithm SHAKE. First, the systems were minimized for 20000 steps using steepest descent to remove steric clashes followed by conjugate gradient. The apolar part of the lipids were melted for 1 ns at 310K in the NVT ensemble to get rid of the crystalline organization of the lipid bilayer. Then, the systems were equilibrated at constant temperature (310K) and constant pressure (1atm) during 10 ns with constraint on protein followed by 10 ns with constraint on backbone only. Finally, the non-constrained runs were performed at constant temperature (310K) and constant pressure (1atm) for 100 ns with a time step of 2 fs. The coordinates of the evolution of each system were backed up every 0.1 ns corresponding to 50000 steps.

### Analysis

The trajectories were analyzed using VMD software *^35^*. The conformational landscape of BoNT/A was monitored using the plugin “RMSD Trajectory Tool” available on VMD. The hydrogen bonds were characterized between 20 and 100 ns with a cutoff of 3 Å and an angle cutoff of 20° between partners (donor and acceptor). The energies of interaction were measured using the tool “Ligand energy Inspector” available in the software Molegro Molecular Viewer (http://molexus.io/molegro-molecular-viewer/). The images were captured using Pymol and Molegro softwares (http://molexus.io/molegro-molecular-viewer/, https://pymol.org/2/).

## Results

In this manuscript, three independent molecular dynamics simulations of 100 ns each were performed involving BoNT/A1 docked with its membrane receptor SV2A singly glycosylated at the position N573 (**figure 1 A**), or the receptor SV2C singly glycosylated at position N559 (SV2C/1g) (**figure 1 B**) or doubly glycosylated at positions N480 and N559 (SV2C/2g) (**figure 1 C**). For each simulation, we monitored the Root Mean Square Deviation (RMSD) of BoNT/A1 to make sure that our solubilized structures do not undergo brutal conformational changes. The plots for the RMSD of Hc-BoNT/A in the system SV2C/1g, SV2C/2g or SV2A are presented in **figure S1 A, B and C respectively.** Their analysis shows that the RMSD transient between 1.5 and 2.5 Å, suggesting that the structures do not go through unrealistic conformational changes generated by a non-stable system.

**Figure 1:**
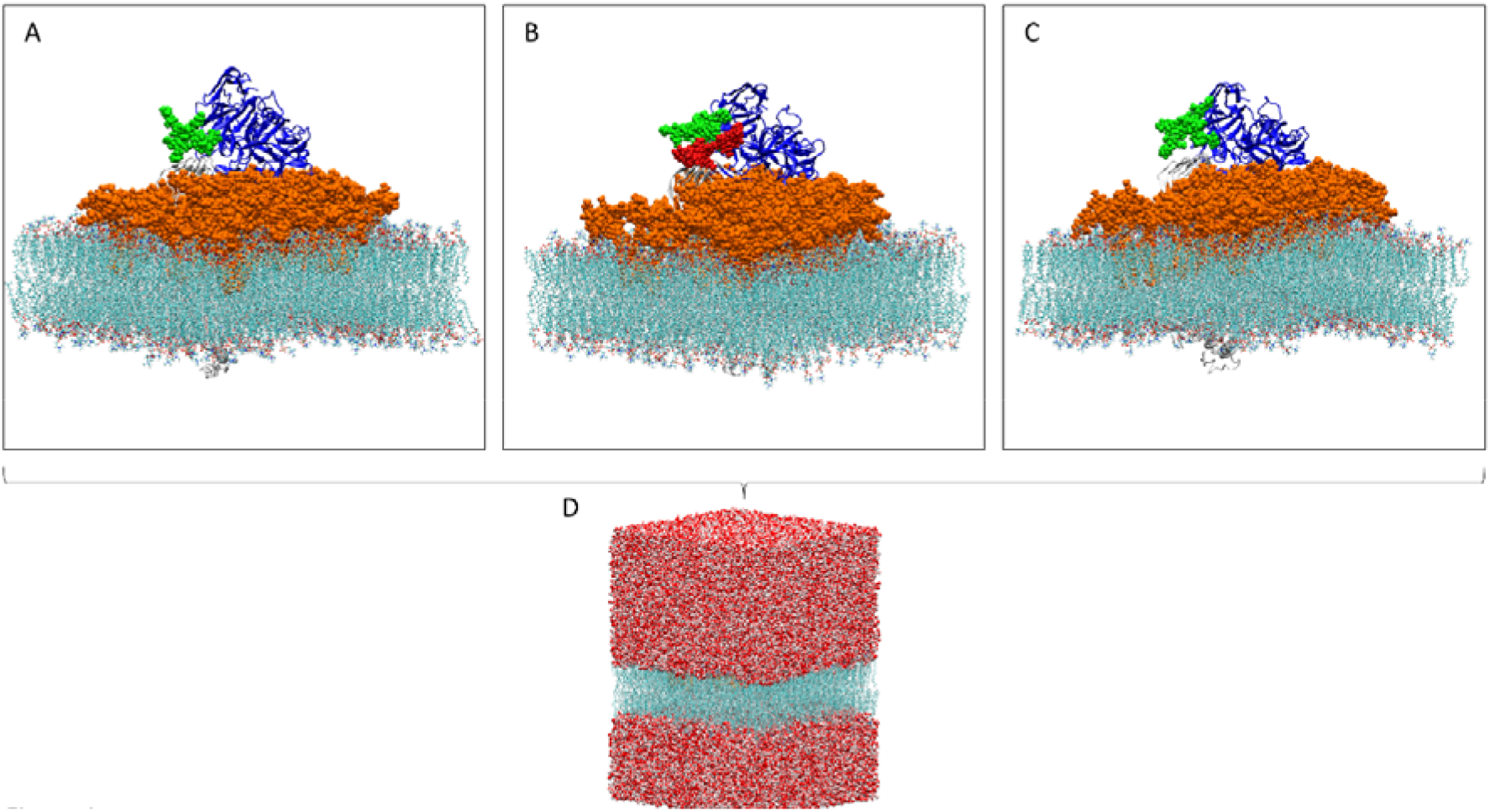
Overview of the BoNT/A1-SV2C1g system **(A)**, BoNT/A1-SV2C/2g **(B)** and BoNT/A1-SV2A **(C)** inserted in a lipid raft environment. The proteins are represented as cartoon colored in blue and white for BoNT/A1 and SV2s respectively. The lipids are represented as spheres colored in orange for the gangliosides and in light salmon for POPC and cholesterol. The glycosylation at position N559 in SV2C or N573 in SV2A are represented as green spheres while the glycosylation at position N480 in SV2C is represented as red spheres. Snapshot of the system after solubilization process **(D)**. Each system was solubilized with a padding greater than 15 Å for the −z and +z axes.

### The glycosylation N480 of human SV2C enhances the contacts of the glycosylation N559 to the surface of BoNT/A1

To make a realistic model of Hc-BoNT/A1 interacting with its membrane receptor, it is of critical importance to consider the N-glycosylation of SV2s. **Figure S2** shows a molecular representation of the full glycosylated human SV2C at positions 480, 484, 534, 559 and 565. The N-glycosylations at positions 480 and 565 (colored in red) are not homologous with the two other isoforms of SV2 while the glycosylations in green (484, 534 and 559) are. Due to its close proximity to the toxin binding site, the glycosylation of LD-SV2C at the position N565 was suspected to interact with Hc-BoNT/A1, giving more stability to the complex SV2C-toxin and leading to improve its affinity with this isoform *^21^*. Nevertheless, the authors concluded that only the glycosylation at position 559 is important for the binding of BoNT/A.

If we consider SV2C with only a glycosylation at the position N559 (N559g), this glycosylation would also interact with residues of LD-SV2C and so it could lead to a loss of contact with BoNT/A1. Due to its opposite position with N559g at the tip of LD-SV2C, we hypothesized that the glycosylation N480 (N480g) could play a role for the binding of BoNT/A1 by preventing N559g to be highly attracted by the amino acids of LD-SV2C. To address this question, we have first plotted the number of hydrogen bonds (H-bonds) established between the glycans and the protein (LD-SV2C and Hc-BoNT/A1) over time (see the part “analysis” of material and methods for the acceptance parameters of the hydrogen bonds quantification). In the system SV2C/2g, N559g established a higher amount of H-bonds with the toxin compared to the system SV2C/1g (**Figure 2 A and B**). Also, more H-bonds were established between LD-SV2C and N559g in the system SV2C/1g compared to the system SV2C/2g (**figure 2 C and D**) and mainly the charged residues Asp, Glu, Arg and Lys are involved as represented by the plot in **figure 2 E**. To complete our analysis, we plotted the H-bonds establishment of N480g with all residues of LD-SV2C and the charged residues Asp, Glu and Lys of LD-SV2C (**Figure 2 F and G**). The data showed that N480g is mainly attracted on the surface of LD-SV2C by charged residues and it establishes more H-bonds with SV2C than N559g. In addition to the analysis of the H-bonds establishment, we measured the total energy of interaction of N559g with the neurotoxin at 40, 60, 80 and 100 ns in order to consider the contribution of every inter-molecular non-covalent bond. The total energy of interaction of N559g in the SV2C/2g system is much higher and stabilized over time than the SV2C/1g system (**Table S1**).

**Figure 2:**
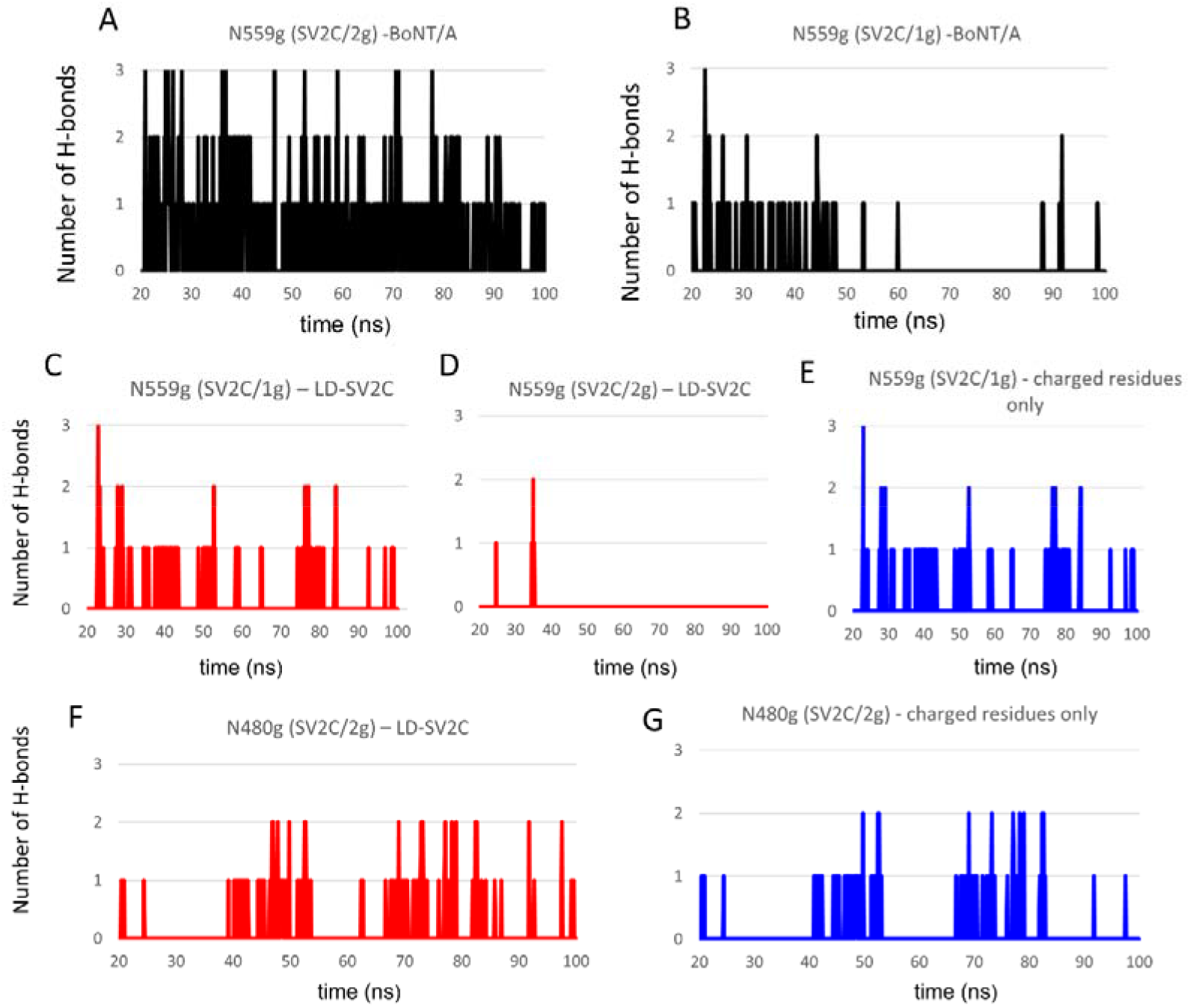
Plot showing the number of H-bonds established between N559g and BoNT/A1 in the system SV2C/2g **(A)**, N559g and BoNT/A1 in the system SV2C/1g **(B)**, N559g and LD-SV2C in the system SV2C/1g **(C)**, N559 and LD-SV2C in the system SV2C/2g **(D)**, N559g and the charged residues in LD-SV2C in the system SV2C/1g **(E)**, N480g and LD-SV2C in the system SV2C/2g **(F)** and N480g and the charged residues of LD-SV2C in the system SV2C/2g.

Together, our computational data suggest that N480g could act as a shield to prevent the attractive long-range interactions exerted by the charged residues of LD-SV2C from focusing their attraction solely on N559g and then resulting in an increase of the binding of N559g to the surface of BoNT/A1.

### Structural analysis of the protein-protein interaction of BoNT/A1-hSV2Cg and BoNT/A1-hSV2Ag complexes

It has been experimentally shown by pull-down experiments that BoNT/A1 has a better affinity for the non-glycosylated rat LD-SV2C than for the non-glycosylated rat LD-SV2A *^19^*. To assess if there are structural differences between BoNT/A1-glycosylated human SV2C (hSV2Cg) and BoNT/A1-glycosylated human SV2A (hSV2Ag) complexes, we compared the structure of each model obtained after 100 ns of simulation. The structural analysis at 100 ns revealed that there is a better complementarity between the beta structure of BoNT/A1 (defined by the sequence 1141-GSVMTT-1146) and hSV2Cg (defined by the sequence 556-EFKNCSFFH-564) (**figure 3 A, B and E**) than BoNT/A1 and hSV2Ag (defined by the sequences 570-RLINSTFLH-578) (**figure 3 C, D and E**). In the case of the BoNT/A1-hSV2Ag complex, we observed an important destructuration of beta strands. Unlike the BoNT/hSV2Cg complex (**figure 3 A, B and E**), the snapshots in **figure 3 C, D and E** show that there is a mismatch of contact between BoNT/A1 and hSV2Ag (highlighted by a circle). Also, the molecular details suggested that this mismatch is predominantly caused by a loose contact of the residues T1145 and T1146 and thus the H-bonds established between these residues and hSV2A (**figure 3 E**). To have a better estimation of the number of H-bonds formed by T1145 and T1146 with hSV2Cg versus hSV2Ag, we quantified the H-bonds establishment of these residues throughout the simulation. The results are plotted in **figure 4 A**, the analysis of the plot clearly shows that more H-bonds are established by T1145 and T1146 with hSV2Cg while only 1 H-bond is established by T1145 and T1146 with the receptor hSV2Ag and no H-bonds were established with this receptor in advanced time between 85 ns and 100 ns. Thus, T1145 and T1146 are more implicated in the binding with hSV2Cg than in the binding with hSV2Ag.

**Figure 3:**
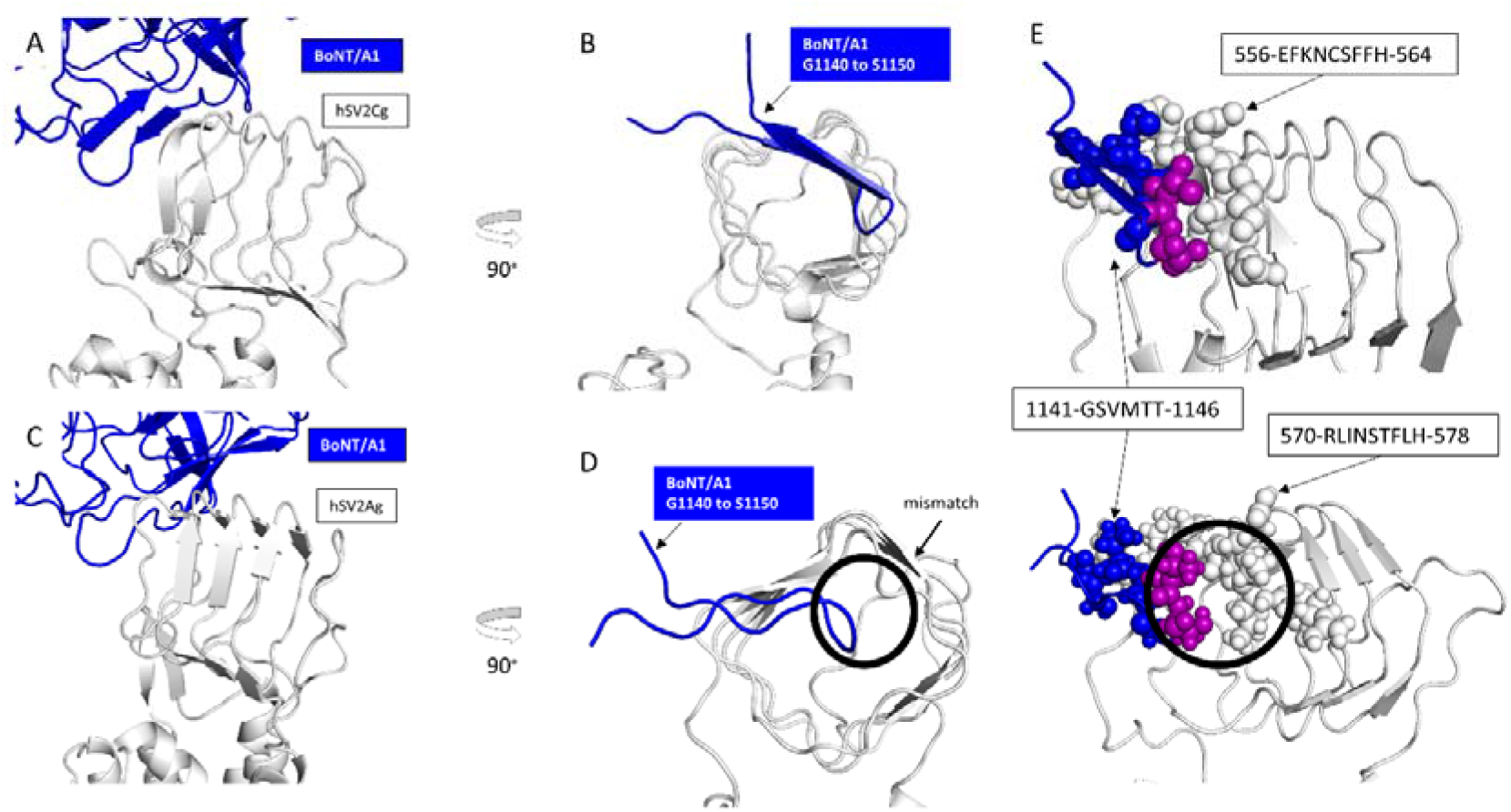
Structure of the complexes BoNT/A1-hSV2Cg **(A and B)** and BoNT/A1-hSV2Ag **(C and D)** obtained after 100 ns of molecular dynamics simulation. Molecular details of the interaction between BoNT/A1-hSV2Cg **(E, top)** or BoNT/A1-hSV2Ag **(E, bottom)**. The amino acids of hSV2C and hSV2A are shown as white spheres while the amino acids of the neurotoxin are shown as blue and purple spheres for the residues T1145 and T1146. The black circle highlights the mismatch between the protein portion of hSV2A and the residues T1145-T1146 of BoNT/A1.

**Figure 4:**
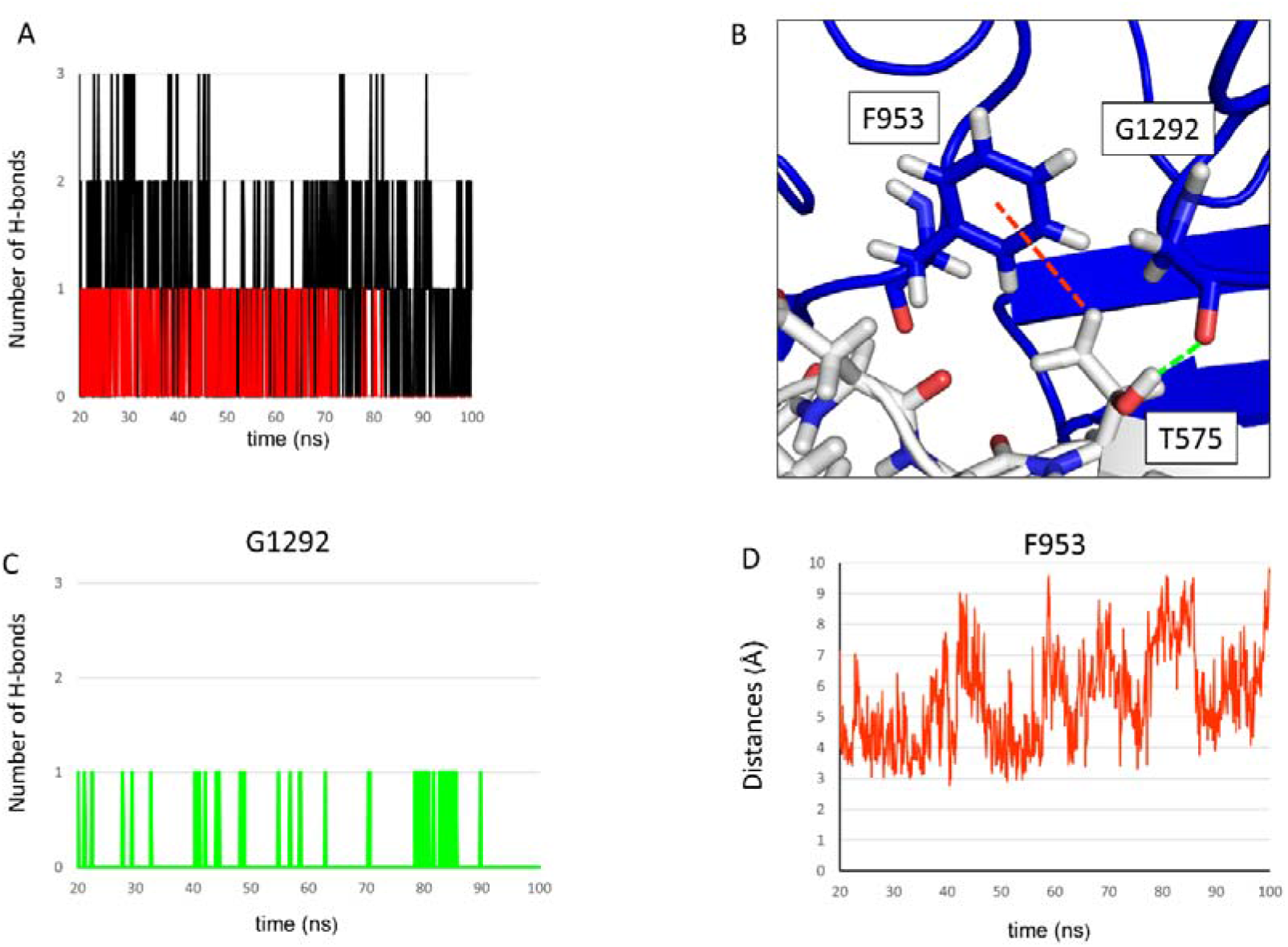
Plot showing the number of H-bonds established by T1145 and T1146 with hSV2Cg (black lines) or hSV2Ag (red lines) **(A)**. Molecular details of the interaction between F953-T575 and G1292-T575 **(B)**. Plot showing the number of H-bonds formed between the backbone of G1292 and T575 **(C)**. Plot of the distances between the lateral aromatic ring of F953 and the CH_3_ group at the lateral chain of T575 **(D)**.

As a response to the mismatch between BoNT/A1 and the protein part of hSV2Ag, we observed that the residues F953 and G1292 interact with the residue T575 of hSV2Ag. The molecular analysis revealed that G1292 interacts with T575 via a H-bond and F953 interacts with the same residue via a CH-π interaction (**figure 4 B**). To evaluate the contribution of these bonds in hSV2Ag binding, we have plotted the H-bonds establishment between G1292 and T575. The plot in **figure 4 C** reveals that G1292 is more in an unbound state with T575 between 20 and 70 ns whilepurp at times between 78 and 87 ns G1292 is highly bound with T575. However, no H-bonds were established between 90 and 100 ns. Next, we plotted the distance between F953 and T575 between 20 and 100 ns. In general, two residues are considered as interacting through a CH-π interaction when the distance is between 2.5 and 5 Å *^38^*. The analysis of the distances plotted in **figure 4 D** show that F953 juggles between the bound and unbound state and F953 is more bound to T575 via CH-π interaction in the first half of the simulation (between 20 to 60 ns) while it is more in an unbound state for the last half of the simulation (60 to 100 ns). So, our results suggest that F953 and G1292 participate in the stabilization by improving the contacts of BoNT/A1 with the protein portion of hSV2Ag by interacting with the residue T575.

In addition to the good surface complementarity between BoNT/A1 and hSV2Cg, we found that the residues P1139 and R1156 enter in interaction with the protein receptor. As shown in **figure S3 A**, P1139 enters in contact with hSV2Cg via a CH-π interaction with H564 while R1156 interacts with the protein receptor via a cation-π interaction *^39^* with F563. We have plotted the distances between F563-R1156 and H564-P1139 over the time to evaluate the durability of these interactions. The plots start at 20 ns and finish at 100 ns. The distances between the NH group of the lateral chain of R1156 and the aromatic ring of F563 are globally under 5 Å, suggesting that the interaction between them is strong and stable over time (**figure S3 B**). For the residues P1139 and H564, the distances reveal that these residues start to enter in contact at 40 ns and the interaction between them is stable over time (**figure S3 C**). These results suggest that the lateral chain of R1156 and P1139 enter durably in interaction with hSV2Cg.

### Interactions of N559g (hSV2C) and N573g (hSV2A) on the surface of BoNT/A1

It is important to consider the glycosylation of SV2A and SV2C in the binding of BoNT/A1 since it has been shown that the glycosylation is critical for the neurotoxin to bind the surface of neurons *^22^*. In addition, mutated BoNT/A1 at positions F953 and G1292 displayed a reduced binding than the wild type for hSV2Cg whereas no difference of binding were observed for non-glycosylated hSV2C, suggesting a different binding strategy adopted by the neurotoxin between these two forms *^22^*. Then, it is of primary importance to consider the glycosylation of SV2 to make a model that predicts the binding of BoNT/A1 with its membrane receptor in a physiological context. At each putative N-glycosylation sites (N573 and N559 for SV2A and SV2C, respectively), we covalently attached a glycosylation that has the same sequence of sugar residues as the glycosylation resolved by MS-spectrometry *^21^*. To explore the mechanism of interaction of N559g and N573g with the neurotoxin, we measured the energy of interaction of each amino acid of BoNT/A1 with the glycosylation at 40, 60, 80 and 100 ns. The results are summarized **in figure S4 and figure S5** for N559g and N573g respectively. Below each table of values, we presented a snapshot showing the global contacts of N559g or N573g with the surface of the toxin at the corresponding timepoint. Residues with a low interaction energy (threshold fixed at −1.5 kJ/mol) were omitted. The data revealed that N559g interacts with more residues of BoNT/A1 than N573g for which F917, F953, N954, T1063 and H1064 are the strongest contact points. Also, the total energy is higher for N559g, meaning a better affinity for the neurotoxin surface.

It is visible from the snapshots showing the global contact of N573g on the surface of BoNT/A1 that N573g also interacts with LD-SV2A. To evaluate the binding frequency of N573g to the surface of LD-SV2A, we plotted the H-bonds establishment between both partners. The analysis of these plots (**figure 6 A and B)** revealed that H-bonds are frequently established between SV2A and N573g over time and all these H-bonds are formed with charged residues (blue lines) and mainly with N553 (purple lines) which is in close contact with N573g as shown in **figure 6 C.** Thus, our results suggest that N573 encounters the same attraction effect as N559g to the SV2C surface in the SV2C1g system which reduces its contacts with the toxin surface. Interestingly, the total energies of interaction of N573g to BoNT/A1 are higher than the total interaction energies of N559g (**table S1 and figure S5**). After a structural analysis of both LD-SV2Ag and LD-SV2Cg, we figured out that SV2C has more charged residues on the tip of the luminal domain accessible by N559g while SV2A has a lower number (**figure 5 C and D**). So, our analysis suggests that N573g is less attracted toward the surface of LD-SV2A due to a lower amount of charged residues on the tip of LD-SV2A. This leads to a less pronounced attraction of N573g toward the surface of SV2A which does not strongly prevent the binding with the neurotoxin like N559g in the system SV2C/1g **(figure 2 B and table S1).**

**Figure 5:**
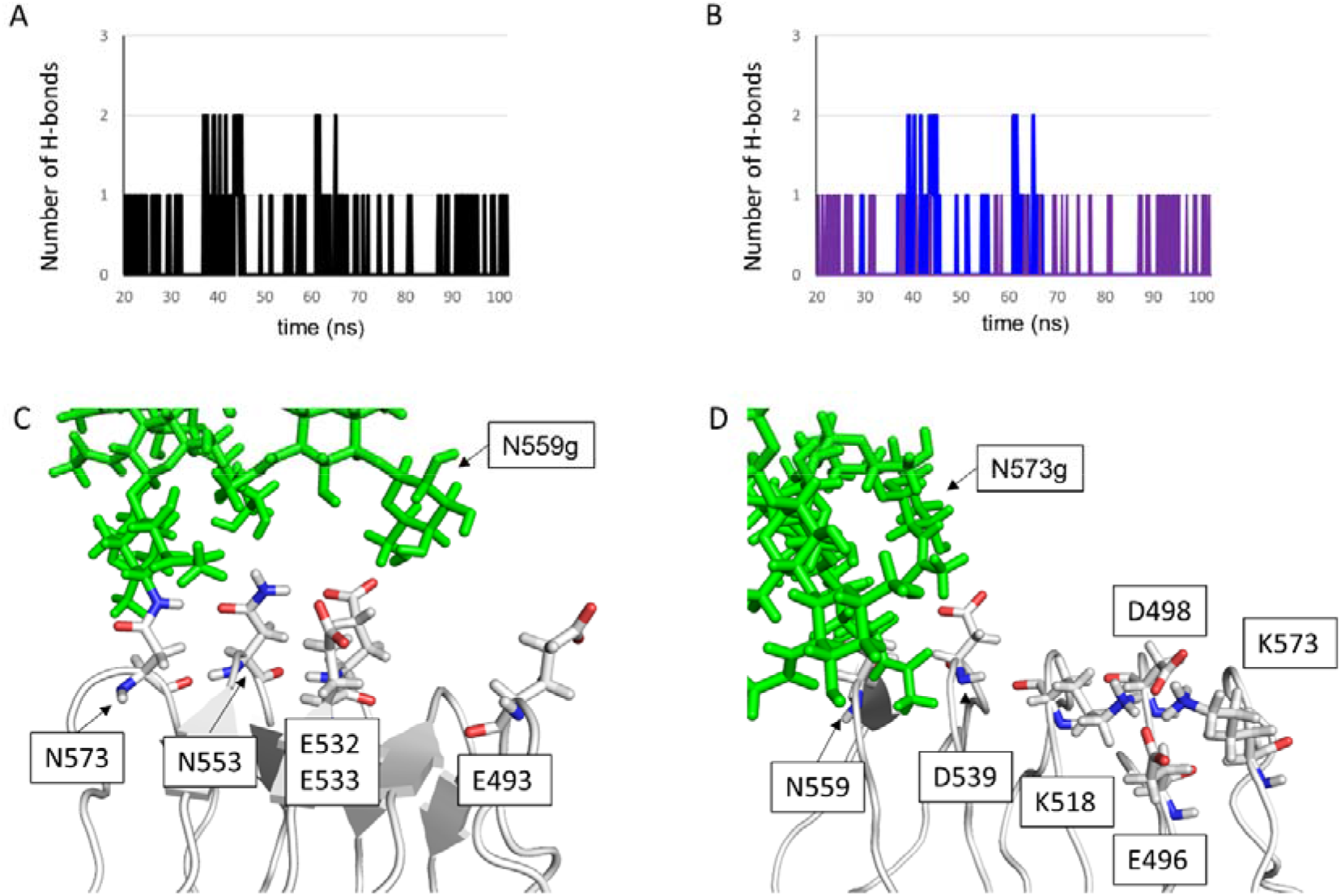
Plot showing the number of H-bonds established by N573g with all residues of LD-SV2A **(A)**. Plot showing the number of H-bonds formed between N573g and all the charged residues of LD-SV2A (blue lines) or the residue N553 (purple lines) **(B)**. Molecular details of LD-SV2A showing the position of N553 and all the charged residues accessible by N573g **(C)**. Molecular details of LD-SV2C showing all the charged residues accessible to N559g **(D)**.

**Figure 6:**
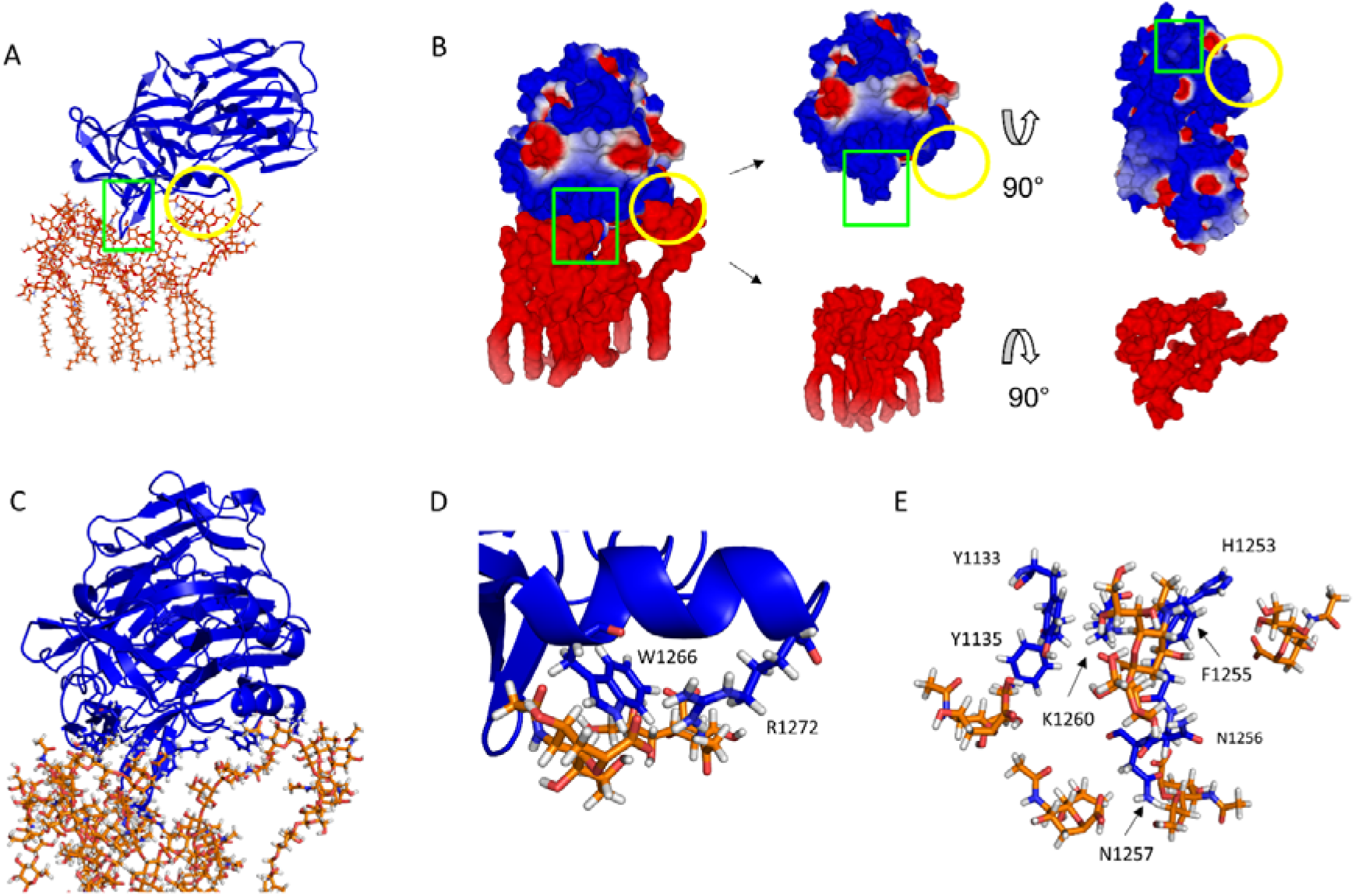
BoNT/A1 interacting with six raft-associated GT1b, the toxin is represented as cartoons colored in blue while the gangliosides are represented as sticks colored in orange **(A)**. Same as previous, but the neurotoxin and the gangliosides are represented as electronic surface in a different orientation (**B**). Molecular details of GT1b interacting in the GBS **(C and D)** or with the GBL **(C and E)**.

### Hc-BoNT/A1 strongly interacts with six lipid raft-associated GT1b

After analyzing the contacts between BoNT/A1 and its glycosylated protein receptors, we investigated the contacts between Hc-BoNT/A1 and the lipid raft-associated GT1b. For this purpose, we analyzed all the contacts between BoNT/A1 and the gangliosides at 100 ns. We found that six GT1b molecules interact via their sialic acids with BoNT/A1 and they particularly target two domains: 1) the structure defined as the gangliosides binding site (GBS), highlighted in a yellow circle in **figure 6 A and B**), 2) the structure we define as the ganglioside binding loop (GBL), highlighted by a green rectangle in the same figure. Interestingly, the GBL is homologous to the Lipid Binding Loop in serotypes B, C, D, D/C and G. To assess the complementarity of the surfaces between the GBL and the lipid raft-associated GT1b, we generated a representation of Hc-BoNT/A1 and the gangliosides in electrostatic surface potential. We can see in **figure 6 B** that the electronic surface of the GBL highlighted by a green square is blue indicating that the global charge in this area is cationic. The electronic surface of the GT1b molecules is red surface meaning that the global charge is anionic due to the presence of sialic acids. Then, the fact that the GBL surface is globally cationic and the GT1b surface is globally anionic, indicate a good chemical complementarity between both partners.

At the hundredth nanosecond, in the GBS, the sialic acids of GT1b molecules were found to mainly interact with amino acids W1266 and R1273 (**figure 6 C and D**). In the GBL region, the GT1b molecules interact with residues H1253, F1255, N1256, N1257 and K1260 (**figure 6 C and E**). In addition to these residues, Y1133 and Y1135 also interact with gangliosides. Asparagine residues mainly interact with the sialic acids of gangliosides via H-bonds. Lysine and arginine side chains could also interact with sialic acid residues via H-bonds in addition to the electrostatic interaction between the cationic group NH_3_^+^ and the anionic group COO^-^. Moreover, the hydroxyl group of tyrosine residues could also establish H-bonds with sialic acids. To evaluate the durability of these H-bonds, we plotted over time the H-bond establishment between the residues Y1133, Y1135, N1256, N1257, K1260 or R1273 and one of the three sialic acids of GT1b. The results revealed that Y1133 and Y1135 (**figure S6 A and B**) do not durably interact with the gangliosides via H-bonds, in contrast with residues N1256, N1257, K1260 and R1273 (**figure S6 C, D, E and F**) suggesting that asparagine, lysine and arginine residues are important contributors of BoNT/A1 binding to the lipid raft.

In addition to the H-bond analysis, we plotted the distances between the sialic acid residues of GT1b and the aromatic residues Y1133, Y1135, H1253 and F1255 of BoNT/A1. The plots revealed that the residues Y1133, H1253 and F1255 interact durably with the sialic acid of the raft-associated GT1b via CH-pi interactions for which F1255 show the most stable interaction followed closely by Y1133 (**figure S7 A, C and D**). The amino acid Y1135 starts to interact with the sialic acid at 85 ns however, its binding is not stable at all and juggles between the bound and unbound state until the end of the trajectory (**figure S7 B**).

## Discussion

In this study, we used molecular dynamics simulations to study the conformational landscape of Hc-BoNT/A1 in complex with its protein receptors hSV2Ag or hSV2Cg. Our strategy was to generate a de novo model instead of relying on Alpha Fold because this algorithm suffers from significant limitations in the case of complex membrane proteins *^40^*. A major outcome of this work is that we considered the full length of hSV2A and hSV2C which is fundamental parameter to insert them into a lipid bilayer and then, it allowed us to simulate the evolution of the complex with respect to the geometric constraints of a neural membrane.

First, we wanted to determine if the N-glycosylation at the position 480 plays a role in the binding of N559g with BoNT/A1. The analysis of the trajectories suggest that N480g improves the surface and the durability of contacts between N559g and the neurotoxin by interacting with the charged residues of LD-SV2C and so, preventing N559g to be attracted toward LD-SV2C as observed in our system SV2C/1g.

Interestingly, we observed that N573g also underwent an attraction to LD-SV2A leading to a loss of surface area and durability of contacts with the BoNT/A1 surface. However, the loss of binding between glycosylation and toxin is less important in SV2A system than in SV2C/1g system, due to a smaller amount of charged residues on the tip of LD-SV2A. Thus, our computational data suggest that BoNT/A1 has a better affinity for the glycosylations on SV2C due to the presence of N480g which is not homologous across SV2s isoforms.

It is well documented in the literature that non-glycosylated SV2C is the receptor which has the highest affinity for BoNT/A1 *^19^*. Our molecular dynamics simulation confirmed this finding by predicting a mismatch of contacts between the beta structure of BoNT/A1 (defined by the sequence 1141-GSVMTT-1146) and SV2A (defined by the sequences 570-RLINSTFLH-578). The molecular details revealed that the lower affinity of BoNT/A1 for SV2A is due to the absence of a CH-π network involving four phenylalanine residues which is present in SV2C. As shown in **figure 7 A**, the CH-π network involving the residues F547, F552, F557 and F562 induces a compaction of the protein portion 556-EFKNCSFFH-564 of LD-SV2C, making its structure accessible by BoNT/A1 via its sequence 1141-GSVMTT-1146. In the case of SV2A, this CH-π network is not present due to a replacement of F557 and F547 in isoform C by two leucine in isoform A. As a result, the structure of SV2A is less compact and therefore more extended (**figure 7 B**). As a result, a mismatch is observed between the residues T1145 and T1146 of the neurotoxin with the surface of SV2A. It was experimentally shown that when T1145 and T1146 are mutated by two alanine, the toxin is no longer able to bind to all SV2 isoforms, highlighting evidence that the mismatch predicted by our simulation can explain the reason for the main difference in affinity between SV2A and SV2C *^20, 22^*. Also, it is worth to note that in our molecular dynamics simulation, the residue F953 of BoNT/A play a more important role for the binding with hSV2Ag by interacting with both the protein portion and the saccharide part of the receptor, while this residue only interact with the glycan part of hSV2Cg **(figure 4, figure S4 and S5)**. Moreover, G1292 appear to only interact with the protein portion of hSV2Ag, while it binds only the glycan portion of hSV2Cg **(figure 4 and figure S4)**. These observations match with the fact that when mutated, the toxin loss its ability to bind SV2A while its binding with SV2C is at least half lost but not totally *^22^*.

**Figure 7:**
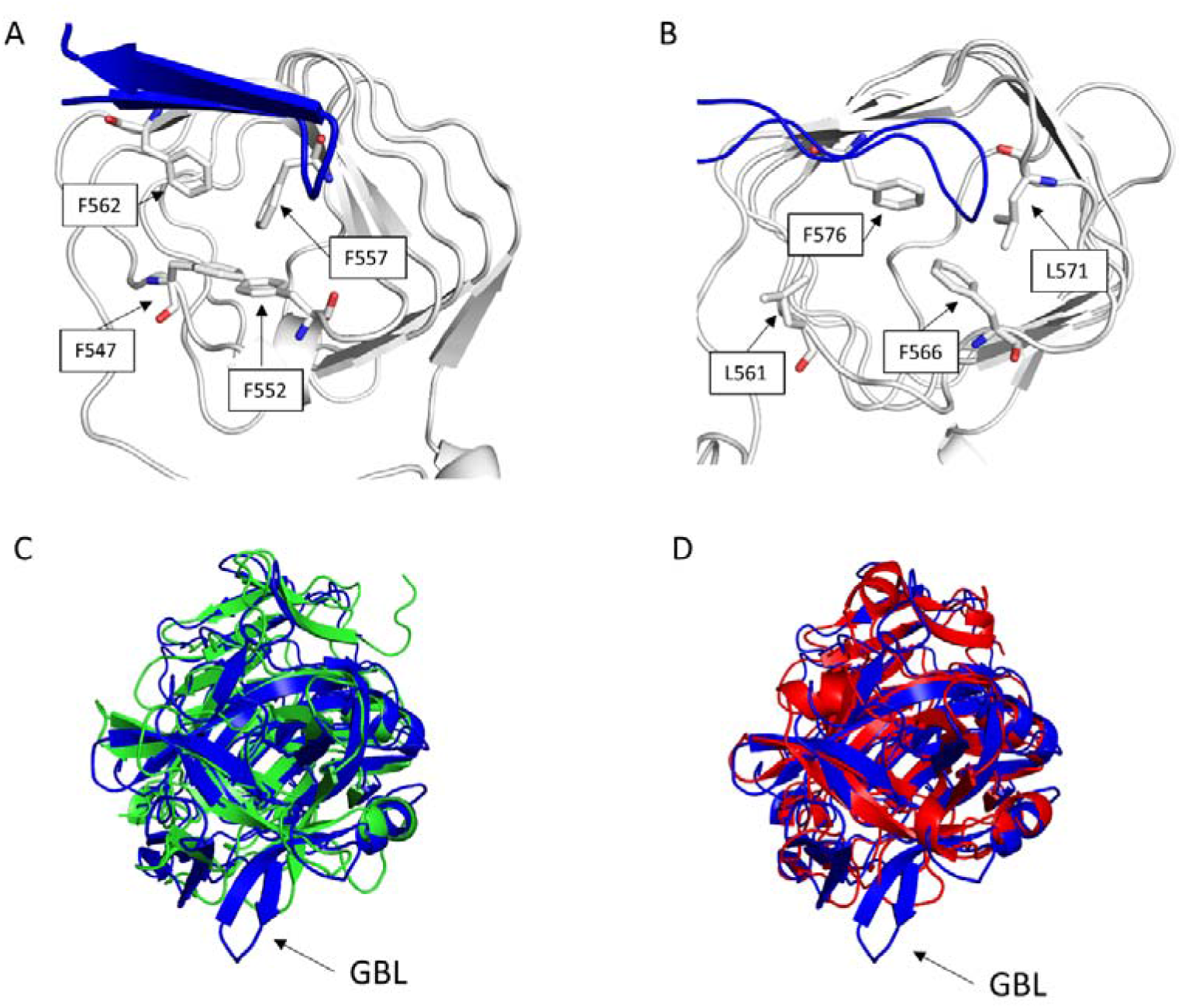
Structural details of LD-SV2C **(A)** and LD-SV2A **(B)**. The four phenylalanine residues causing the compaction of LD-SV2C are shown as white sticks. Structural alignment of BoNT/A1 with BoNT/E (PDB: 3FFZ) **(C)** or with BoNT/F (PDB: 3FUQ) **(D)**. The neurotoxins are represented as cartoon colored in blue for BoNT/A1, green for BoNT/E and red for BoNT/F.

Then, we compared the dynamics of interactions between N559g and N573g with BoNT/A1 at different time in the simulations. In our simulation, in addition to recover the experimentally validated major contact points which are F953, H1064 and G1292, the interaction with those residues is maintained throughout the simulation *^22^*. Also, we observed additional point of contacts for which F917 displayed the strongest energy of interaction and appear in both hSV2Ag and hSV2Cg (**figure S4 and S5).** These secondary interactions allowed a fine tuning of the whole complex.

Globally, the amino acids of BoNT/A1 involved in protein-protein or protein-glycan interactions are not the same in function of the receptor bound by the toxin (here SV2A or SV2C) due to the differences of the surface topology of these receptors. Then, our *In Silico* data suggest a different mechanism of interaction of the neurotoxin with hSV2Ag or hSV2Cg.

During the simulation, we found that in addition to GBS, gangliosides also interact with BoNT/A1 via a structure that we referred to as the Ganglioside Binding Loop “GBL”, referring to the appellation used in previous studies *^41, 42^*. The GBL is defined by the amino acid sequence 1253-HQFNNIAK-1260 for which H1253 was identified to interact with ganglioside *^43^*. After performing a structural alignment of the heavy chain of BoNT/A1, BoNT/E1 and BoNT/F1, we observed that the GBL is not conserved in serotype E1 and F1 (**figure 7 C and D**). The absence of GBL in BoNT/E1 and BoNT/F1 may be one of the reasons why BoNT/A1 has a lower LD50 in mice treated with BoNTs by intraperitoneal injection *^44^*.

To summarize, our molecular dynamics simulations suggest a global model in which BoNT/A1 interact with its membrane receptor complex which includes i) the glycosylated luminal domain of SV2C, and ii) six molecules of GT1b which interact with the HcN subdomain of BoNT/A1 via its GBS and its GBL structure (**Figure 8**).

**Figure 8:**
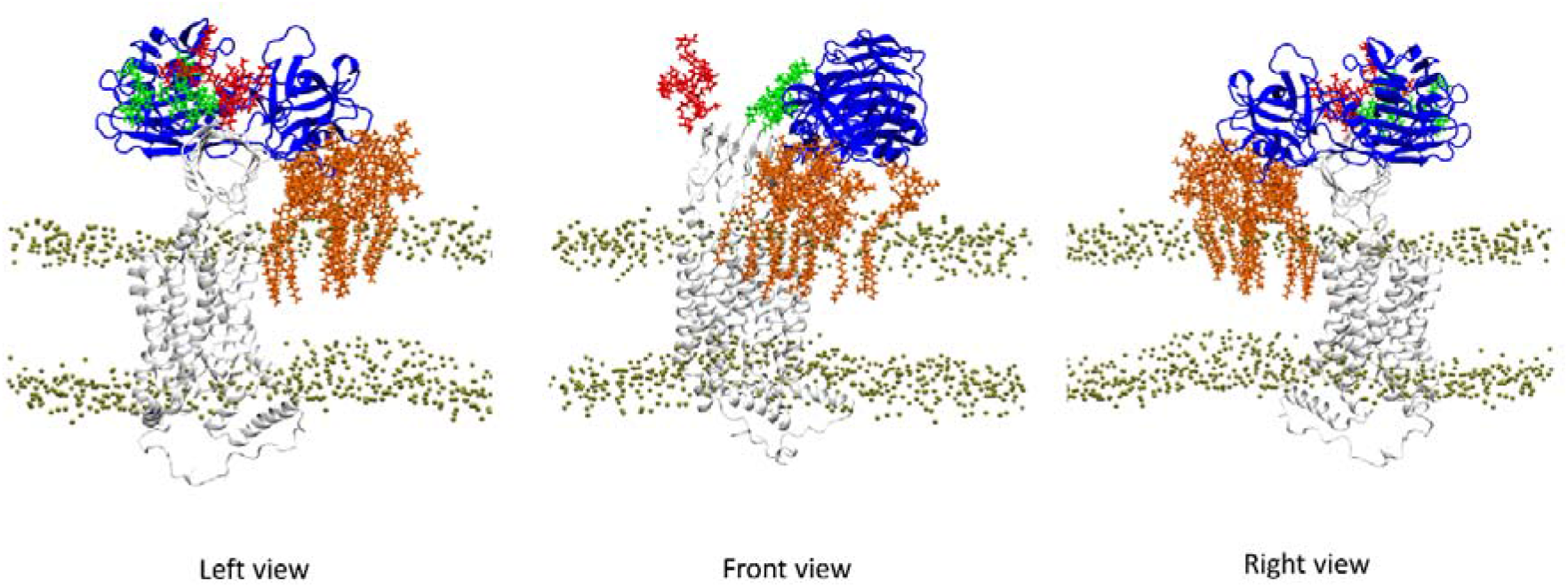
Model of BoNT/A1 interacting with its membrane receptor hSV2C and six molecules of GT1b (obtained after 100 ns of simulation).

Overall, our data give new insights on the molecular mechanisms controlling the binding of botulinum neurotoxin A1 to SV2 receptors. We solved the puzzle generated by mutational studies that could be only partially understood with crystallographic data lacking both a biologically relevant membrane environment and a full glycosylation of SV2. Considering that this toxin is the most potent serotype in humans, our molecular model could also be useful to design specific inhibitors directed against this threat.

## Supporting information

RMSD, molecular model, atom distances, energy of interactions

## References

[1] Andersson, A., Rönner, U., and Granum, P. E. (1995) What problems does the food industry have with the spore-forming pathogens Bacillus cereus and Clostridium perfringens?, International Journal of Food Microbiology 28, 145–155.

[2] Vinh, D. C. (2005) Strains and toxins of &lt;em&gt;Clostridium&lt;/em&gt, Canadian Medical Association Journal 172, 312.

[3] Dong, M., Masuyer, G., and Stenmark, P. (2019) Botulinum and Tetanus Neurotoxins, Annual Review of Biochemistry 88, 811–837.

[4] Hörman, A., Nevas, M., Lindström, M., Hänninen, M. L., and Korkeala, H. (2005) Elimination of botulinum neurotoxin (BoNT) type B from drinking water by small-scale (personal-use) water purification devices and detection of BoNT in water samples, Applied and environmental microbiology 71, 1941–1945.

[5] Dong, M., and Stenmark, P. (2021) The Structure and Classification of Botulinum Toxins, In Botulinum Toxin Therapy (Whitcup, S. M., and Hallett, M., Eds.), pp 11–33, Springer International Publishing, Cham.

[6] Fields, R. D. (1998) Clostridial Neurotoxins in Synaptic Research, The Neuroscientist: a review journal bringing neurobiology, neurology and psychiatry 4, 324–328.

[7] Zhang, S., Masuyer, G., Zhang, J., Shen, Y., Lundin, D., Henriksson, L., Miyashita, S.-L, Martínez-Carranza, M., Dong, M., and Stenmark, P. (2017) Identification and characterization of a novel botulinum neurotoxin, Nature Communications 8, 14130.

[8] Sen, E., Kota, K. P., Panchal, R. G., Bavari, S., and Kiris, E. (2021) Screening of a Focused Ubiquitin-Proteasome Pathway Inhibitor Library Identifies Small Molecules as Novel Modulators of Botulinum Neurotoxin Type A Toxicity, Frontiers in pharmacology 12, 763950.

[9] Maslanka, S. E. (2014) Botulism as a Disease of Humans, In Molecular Aspects of Botulinum Neurotoxin (Foster, K. A., Ed.), pp 259–289, Springer New York, New York, NY.

[10] Verderio, C., Rossetto, O., Grumelli, C., Frassoni, C., Montecucco, C., and Matteoli, M. (2006) Entering neurons: botulinum toxins and synaptic vesicle recycling, EMBO reports 7, 995–999.

[11] Nigam, P., and Nigam, A. (2010) Botulinum toxin, Indian Journal of Dermatology 55, 8–14.

[12] Fu, Z., Huang, H., and Huang, J. (2022) Efficacy and safety of botulinum toxin type A for postoperative scar prevention and wound healing improvement: A systematic review and meta-analysis, Journal of cosmetic dermatology 21, 176–190.

[13] Zakin, E., and Simpson, D. M. (2021) Botulinum Toxin Therapy in Writer&rsquo;s Cramp and Musician&rsquo;s Dystonia, Toxins 13, 899.

[14] An, Y., Kim, Y.-J., Kim, C.-s., Yim, H., Kim, M., Lee, E.-K., Oh, H.-J., Han, J.-H., Yoo, E., Kim, S., Woo, J., Moore, E. R. B., Jung, J.-Y., and Park, W. (2021) Therapeutic efficacy of new botulinum toxin identified in CCUG 7968 strain, Applied Microbiology and Biotechnology 105, 8727–8737.

[15] Pirazzini, M., Rossetto, O., Eleopra, R., and Montecucco, C. (2017) Botulinum Neurotoxins: Biology, Pharmacology, and Toxicology, Pharmacological Reviews 69, 200–235.

[16] Rossetto, O., Pirazzini, M., and Montecucco, C. (2014) Botulinum neurotoxins: genetic, structural and mechanistic insights, Nature Reviews Microbiology 12, 535–549.

[17] Kim, D.-W., Lee, S.-K., and Ahnn, J. (2015) Botulinum Toxin as a Pain Killer: Players and Actions in Antinociception, Toxins 7, 2435–2453.

[18] Dong, M., Yeh, F., Tepp, W. H., Dean, C., Johnson, E. A., Janz, R., and Chapman, E. R. (2006) SV2 Is the Protein Receptor for Botulinum Neurotoxin A, Science 312, 592–596.

[19] Weisemann, J., Stern, D., Mahrhold, S., Dorner, B. G., and Rummel, A. (2016) Botulinum Neurotoxin Serotype A Recognizes Its Protein Receptor SV2 by a Different Mechanism than Botulinum Neurotoxin B Synaptotagmin, Toxins 8, 154.

[20] Benoit, R. M., Frey, D., Hilbert, M., Kevenaar, J. T., Wieser, M. M., Stirnimann, C. U., McMillan, D., Ceska, T., Lebon, F., Jaussi, R., Steinmetz, M. O., Schertler, G. F. X., Hoogenraad, C. C., Capitani, G., and Kammerer, R. A. (2014) Structural basis for recognition of synaptic vesicle protein 2C by botulinum neurotoxin A, Nature 505, 108–111.

[21] Mahrhold, S., Bergström, T., Stern, D., Dorner, B. G., Åstot, C., and Rummel, A. (2016) Only the complex N559-glycan in the synaptic vesicle glycoprotein 2C mediates high affinity binding to botulinum neurotoxin serotype A1, The Biochemical journal 473, 2645–2654.

[22] Yao, G., Zhang, S., Mahrhold, S., Lam, K.-h., Stern, D., Bagramyan, K., Perry, K., Kalkum, M., Rummel, A., Dong, M., and Jin, R. (2016) N-linked glycosylation of SV2 is required for binding and uptake of botulinum neurotoxin A, Nature Structural & Molecular Biology 23, 656–662.

[23] Dong, M., Liu, H., Tepp, W. H., Johnson, E. A., Janz, R., and Chapman, E. R. (2008) Glycosylated SV2A and SV2B Mediate the Entry of Botulinum Neurotoxin E into Neurons, Molecular Biology of the Cell 19, 5226–5237.

[24] Rummel, A., Häfner, K., Mahrhold, S., Darashchonak, N., Holt, M., Jahn, R., Beermann, S., Karnath, T., Bigalke, H., and Binz, T. (2009) Botulinum neurotoxins C, E and F bind gangliosides via a conserved binding site prior to stimulation-dependent uptake with botulinum neurotoxin F utilising the three isoforms of SV2 as second receptor, Journal of Neurochemistry 110, 1942–1954.

[25] Azzaz, F., Yahi, N., Di Scala, C., Chahinian, H., and Fantini, J. (2022) Chapter Eight - Ganglioside binding domains in proteins: Physiological and pathological mechanisms, In Advances in Protein Chemistry and Structural Biology (Donev, R., Ed.), pp 289–324, Academic Press.

[26] Fantini, J., Yahi, N., Azzaz, F., and Chahinian, H. (2021) Structural dynamics of SARS-CoV-2 variants: A health monitoring strategy for anticipating Covid-19 outbreaks, The Journal of infection 83, 197–206.

[27] Yahi, N., and Fantini, J. (2014) Deciphering the glycolipid code of Alzheimer’s and Parkinson’s amyloid proteins allowed the creation of a universal ganglioside-binding peptide, PloS one 9, e104751.

[28] Fantini, J., Garmy, N., Mahfoud, R., and Yahi, N. (2004) Lipid rafts: structure, function and role in HIV, Alzheimer’s and prion diseases, Expert Reviews in Molecular Medicine 4, 1–22.

[29] Yahi, N., Di Scala, C., Chahinian, H., and Fantini, J. (2022) Innovative treatment targeting gangliosides aimed at blocking the formation of neurotoxic α-synuclein oligomers in Parkinson’s disease, Glycoconjugate Journal 39, 1–11.

[30] Froimowitz, M. (1993) HyperChem: a software package for computational chemistry and molecular modeling, BioTechniques 14, 1010–1013.

[31] Wu, E. L., Cheng, X., Jo, S., Rui, H., Song, K. C., Dávila-Contreras, E. M., Qi, Y., Lee, J., Monje-Galvan, V., Venable, R. M., Klauda, J. B., and Im, W. (2014) CHARMM-GUI Membrane Builder toward realistic biological membrane simulations, Journal of Computational Chemistry 35, 1997–2004.

[32] Lee, J., Patel, D. S., Ståhle, J., Park, S. J., Kern, N. R., Kim, S., Lee, J., Cheng, X., Valvano, M. A., Holst, O., Knirel, Y. A., Qi, Y., Jo, S., Klauda, J. B., Widmalm, G., and Im, W. (2019) CHARMM-GUI Membrane Builder for Complex Biological Membrane Simulations with Glycolipids and Lipoglycans, Journal of chemical theory and computation 15, 775–786.

[33] Jo, S., Kim, T., Iyer, V. G., and Im, W. (2008) CHARMM-GUI: A web-based graphical user interface for CHARMM, Journal of Computational Chemistry 29, 1859–1865.

[34] Jo, S., Cheng, X., Islam, S. M., Huang, L., Rui, H., Zhu, A., Lee, H. S., Qi, Y., Han, W., Vanommeslaeghe, K., MacKerell, A. D., Roux, B., and Im, W. (2014) Chapter Eight - CHARMM-GUI PDB Manipulator for Advanced Modeling and Simulations of Proteins Containing Nonstandard Residues, In Advances in Protein Chemistry and Structural Biology (Karabencheva-Christova, T., Ed.), pp 235–265, Academic Press.

[35] Humphrey, W., Dalke, A., and Schulten, K. (1996) VMD: Visual molecular dynamics, Journal of Molecular Graphics 14, 33–38.

[36] Phillips, J. C., Hardy, D. J., Maia, J. D. C., Stone, J. E., Ribeiro, J. V., Bernardi, R. C., Buch, R., Fiorin, G., Hénin, J., Jiang, W., McGreevy, R., Melo, M. C. R., Radak, B. K., Skeel, R. D., Singharoy, A., Wang, Y., Roux, B., Aksimentiev, A., Luthey-Schulten, Z., Kalé, L. V., Schulten, K., Chipot, C., and Tajkhorshid, E. (2020) Scalable molecular dynamics on CPU and GPU architectures with NAMD, The Journal of chemical physics 153, 044130.

[37] Huang, J., Rauscher, S., Nawrocki, G., Ran, T., Feig, M., de Groot, B. L., Grubmüller, H., and MacKerell, A. D. (2017) CHARMM36m: an improved force field for folded and intrinsically disordered proteins, Nature Methods 14, 71–73.

[38] Nishio, M., Umezawa, Y., Fantini, J., Weiss, M. S., and Chakrabarti, P. (2014) CH-π hydrogen bonds in biological macromolecules, Physical chemistry chemical physics: PCCP 16, 12648–12683.

[39] Infield, D. T., Rasouli, A., Galles, G. D., Chipot, C., Tajkhorshid, E., and Ahern, C. A. (2021) Cation-π Interactions and their Functional Roles in Membrane Proteins, Journal of molecular biology 433, 167035.

[40] Azzaz, F., and Fantini, J. (2022) The epigenetic dimension of protein structure, Biomolecular concepts 13, 55–60.

[41] Karalewitz, A. P. A., Kroken, A. R., Fu, Z., Baldwin, M. R., Kim, J.-J. P., and Barbieri, J. T. (2010) Identification of a Unique Ganglioside Binding Loop within Botulinum Neurotoxins C and D-SA, Biochemistry 49, 8117–8126.

[42] Kroken, A. R., Karalewitz, A. P., Fu, Z., Kim, J. J., and Barbieri, J. T. (2011) Novel ganglioside-mediated entry of botulinum neurotoxin serotype D into neurons, The Journal of biological chemistry 286, 26828–26837.

[43] Gregory, K. S., Mojanaga, O. O., Liu, S. M., and Acharya, K. R. (2022) Crystal Structures of Botulinum Neurotoxin Subtypes A4 and A5 Cell Binding Domains in Complex with Receptor Ganglioside, Toxins 14, 129.

[44] Rossetto, O., and Montecucco, C. (2019) Tables of Toxicity of Botulinum and Tetanus Neurotoxins, Toxins 11, 686.

