## Supplementary material for "Structural basis of botulinum neurotoxin serotype A1 binding to human SV2A or SV2C receptors": RMSD, molecular model, atom distances, energy of interactions

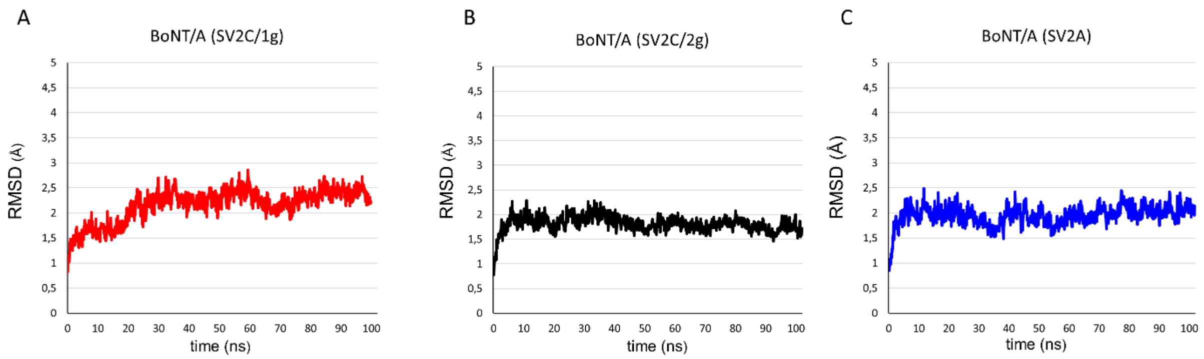


**Figure S1**: RMSD of BoNT/A1 in the system SV2C/1g (red curve) **(A)**, SV2C/2g (black curve) **(B)** and SV2A (blue curve) **(C)**.

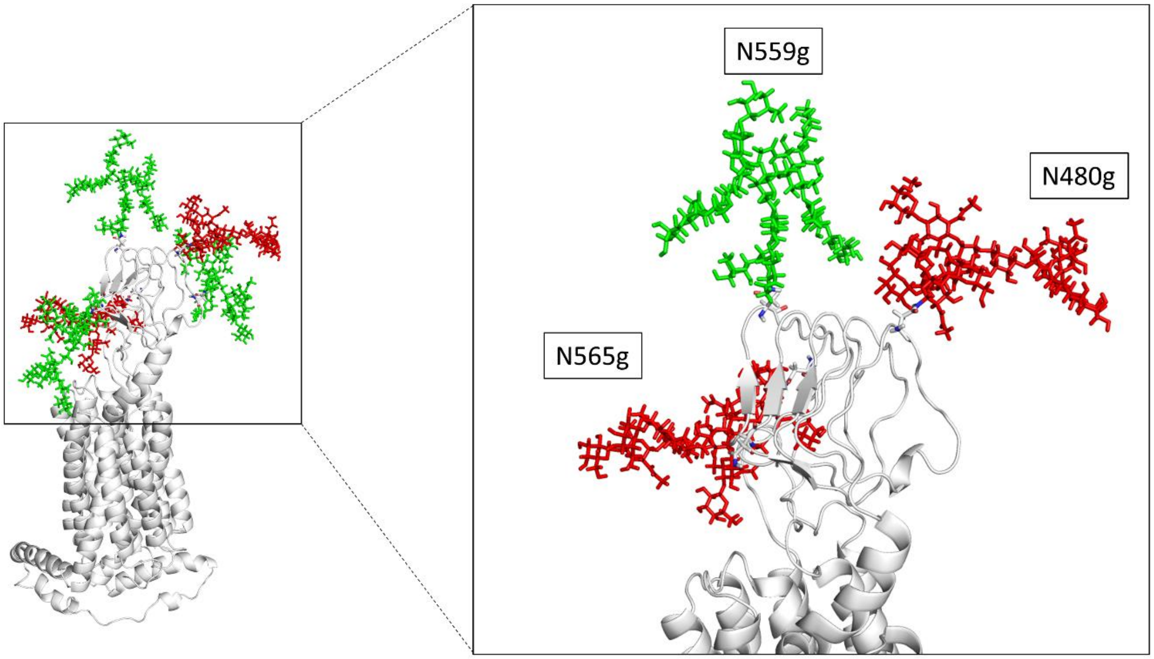


**Figure S2:** Molecular model of SV2C with its five glycans covalently attached at the N-glycosylation putative sites N480, N484, N534, N559 and N565. The glycosylations which are homologous across SV2s isoforms are colored in green while the glycosylations which are specific to SV2C are colored in red. Focus on the glycans at the position N480, N559 and N565. N480g is on the tip of LD-SV2C at the opposite position of N559g. This glycosylation could play a role in improving the contacts of N559g on the surface of BoNT/A1.

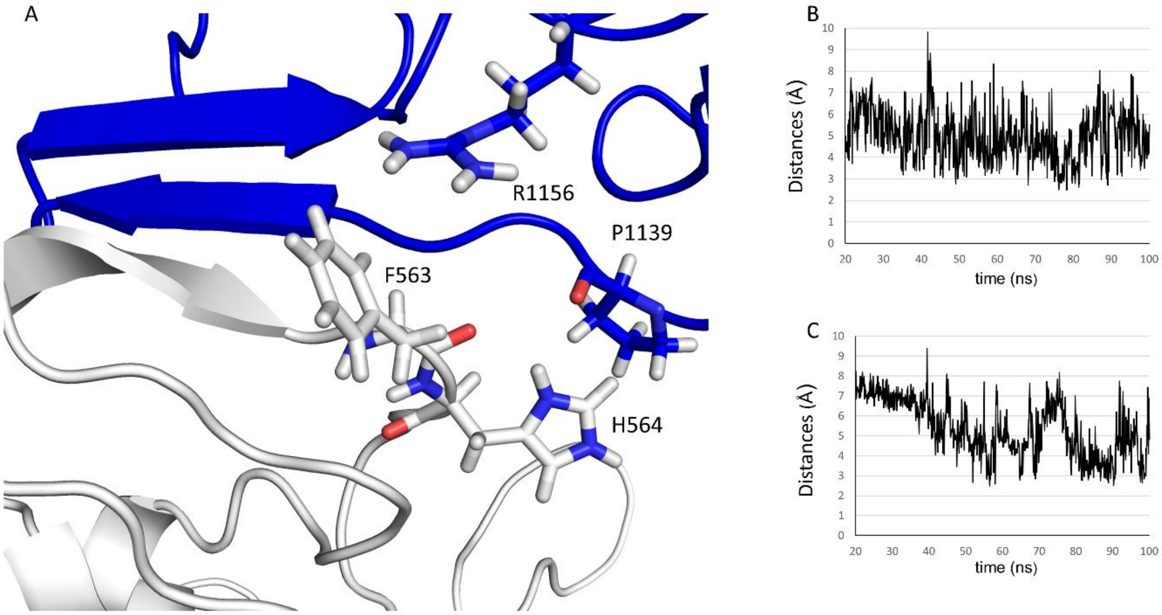


**Figure S3:** Molecular details of the interaction between the lateral chain of R1156 with the ring of F563 or the lateral chain of P1139 and the ring of H564 **(A)**. Plot showing the distance between the lateral chain of R1156 and the aromatic ring of F563 **(B)**, plot showing the distances between P1139 and H564 **(C)**.

| Amino acids | Position in the protein sequence | Energy of interaction (kJ/mol) |
|---|---|---|
| Arg | 1061 | -17.386 |
| Asn | 954 | -15.3371 |
| Asp | 1062 | -2.59897 |
| Asp | 1289 | -14.8752 |
| Cys | 1060 | -5.96747 |
| Gly | 1290 | -3.79047 |
| Gly | 1292 | -3.37452 |
| His | 1064 | -13.3854 |
| Ile | 956 | -6.02254 |
| Leu | 919 | -2.98994 |
| Lys | 951 | -2.94765 |
| Phe | 953 | -22.9418 |
| Thr | 1063 | -6.81633 |
| Trp | 1291 | -3.26263 |
| | Total : | -121.69602 |

| Amino acids | Position in the protein sequence | Energy of interaction (kJ/mol) |
|---|---|---|
| Arg | 1061 | -6.72515 |
| Asn | 954 | -12.3481 |
| Asp | 1062 | -1.65641 |
| Gly | 1290 | -4.8582 |
| Gly | 1292 | -4.0904 |
| His | 1064 | -17.8699 |
| Ile | 956 | -6.12282 |
| Leu | 919 | -4.63067 |
| Lys | 903 | -9.1641 |
| Lys | 951 | -2.06837 |
| Phe | 917 | -12.4311 |
| Phe | 953 | -19.5369 |
| Ser | 955 | -2.09801 |
| Thr | 1063 | -18.0603 |
| Trp | 1291 | -2.61441 |
| | Total : | -124.27484 |

| Amino acids | Position in the protein sequence | Energy of interaction (kJ/mol) |
|---|---|---|
| Arg | 1061 | -4.4352 |
| Arg | 1294 | -5.02673 |
| Asn | 954 | -15.0135 |
| Gly | 1292 | -4.86002 |
| His | 1064 | -21.4453 |
| Leu | 919 | -3.14569 |
| Lys | 903 | -3.09619 |
| Lys | 951 | -2.31815 |
| Phe | 917 | -17.3114 |
| Phe | 953 | -22.4319 |
| Thr | 1063 | -16.5039 |
| | Total : | -115.58798 |

| Amino acids | Position in the protein sequence | Energy of interaction (kJ/mol) |
|---|---|---|
| Arg | 1294 | -1.87281 |
| Asn | 905 | -2.20672 |
| Asn | 954 | -12.081 |
| Gly | 1290 | -2.09092 |
| Gly | 1292 | -3.37143 |
| His | 1064 | -25.4403 |
| Lys | 903 | -2.2277 |
| Lys | 951 | -2.04409 |
| Phe | 917 | -15.3149 |
| Phe | 953 | -20.2787 |
| Thr | 1063 | -14.5317 |
| Trp | 1291 | -4.8127 |
| | Total : | -106.27297 |

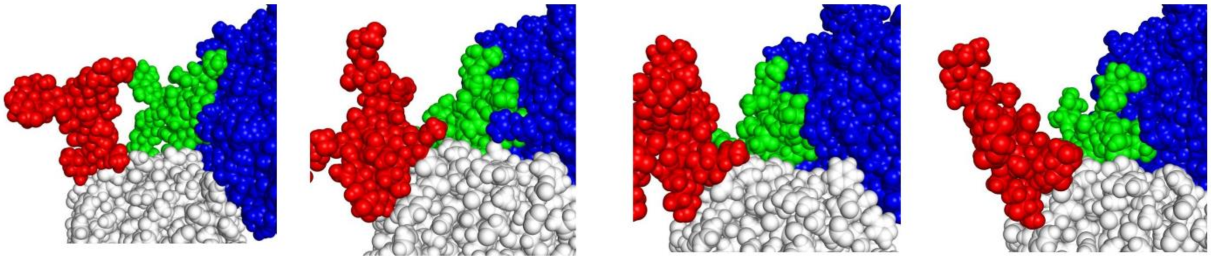

**Figure S4:** Energy of interaction of BoNT/A1 residues for N559g. Below each table of values is associated a snapshot showing the global contacts of the glycan on the surface of BoNT/A1. All elements are represented as spheres: BoNT/A1 is colored in blue, SV2C in white, N559g in green, and N480g in red.

| Amino acids | Position in the protein sequence | Energy of interaction (kJ/mol) |
|---|---|---|
| Asn | 905 | -1.54835 |
| Asp | 1289 | -2.51141 |
| His | 1064 | -23.6699 |
| Lys | 903 | -2.54893 |
| Phe | 953 | -5.20747 |
| Pro | 908 | -3.25898 |
| Pro | 950 | -1.93912 |
| Thr | 1063 | -2.94918 |
| Tyr | 1066 | -14.7552 |
| | Total : | -58.38854 |

| Amino acids | Position in the protein sequence | Energy of interaction (kJ/mol) |
|---|---|---|
| Arg | 1294 | -1.90155 |
| Asn | 905 | -20.833 |
| Asp | 1289 | -12.4185 |
| His | 1064 | -29.1665 |
| Phe | 906 | -2.96035 |
| Phe | 917 | -3.12392 |
| Phe | 953 | -5.12043 |
| Pro | 908 | -5.60878 |
| Ser | 902 | -6.97432 |
| Tyr | 1066 | -2.22393 |
| Val | 904 | -7.65964 |
| | Total : | -97.99092 |

| Amino acids | Position in the protein sequence | Energy of interaction (kJ/mol) |
|---|---|---|
| Asn | 905 | -13.5368 |
| Asp | 1289 | -9.37546 |
| Gln | 915 | -1.66084 |
| His | 1064 | -14.1593 |
| Lys | 903 | -3.17539 |
| Phe | 917 | -4.07793 |
| Phe | 953 | -3.24582 |
| Ser | 902 | -4.18933 |
| Tyr | 1066 | -2.32256 |
| Val | 904 | -2.92314 |
| | Total : | -58.66657 |

| Amino acids | Position in the protein sequence | Energy of interaction (kJ/mol) |
|---|---|---|
| Asn | 905 | -14.6518 |
| His | 1064 | -22.5847 |
| Lys | 903 | -9.34634 |
| Phe | 917 | -16.7081 |
| Phe | 953 | -18.861 |
| Ser | 902 | -6.1647 |
| Tyr | 1066 | -1.63036 |
| Val | 904 | -1.58267 |
| | Total : | -91.52967 |

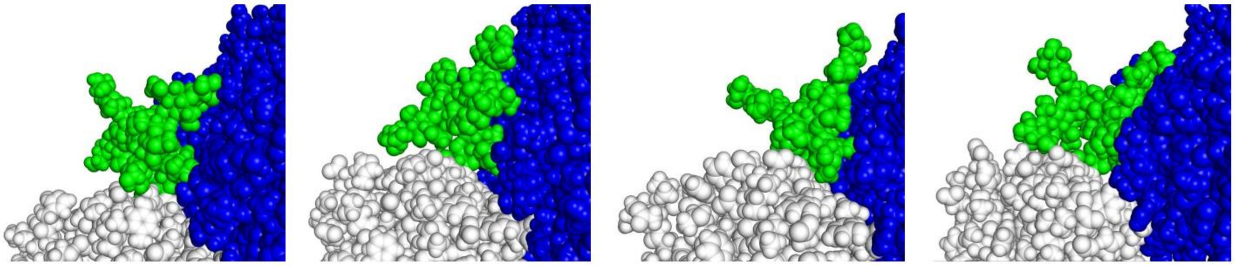

**Figure S5:** Energy of interaction of BoNT/A1 residues for N573g. Below each table of values is associated a snapshot showing the global contacts of the glycan on the surface of BoNT/A1. All elements are represented as spheres: BoNT/A1 is colored in blue, SV2A in white and N579g in green.

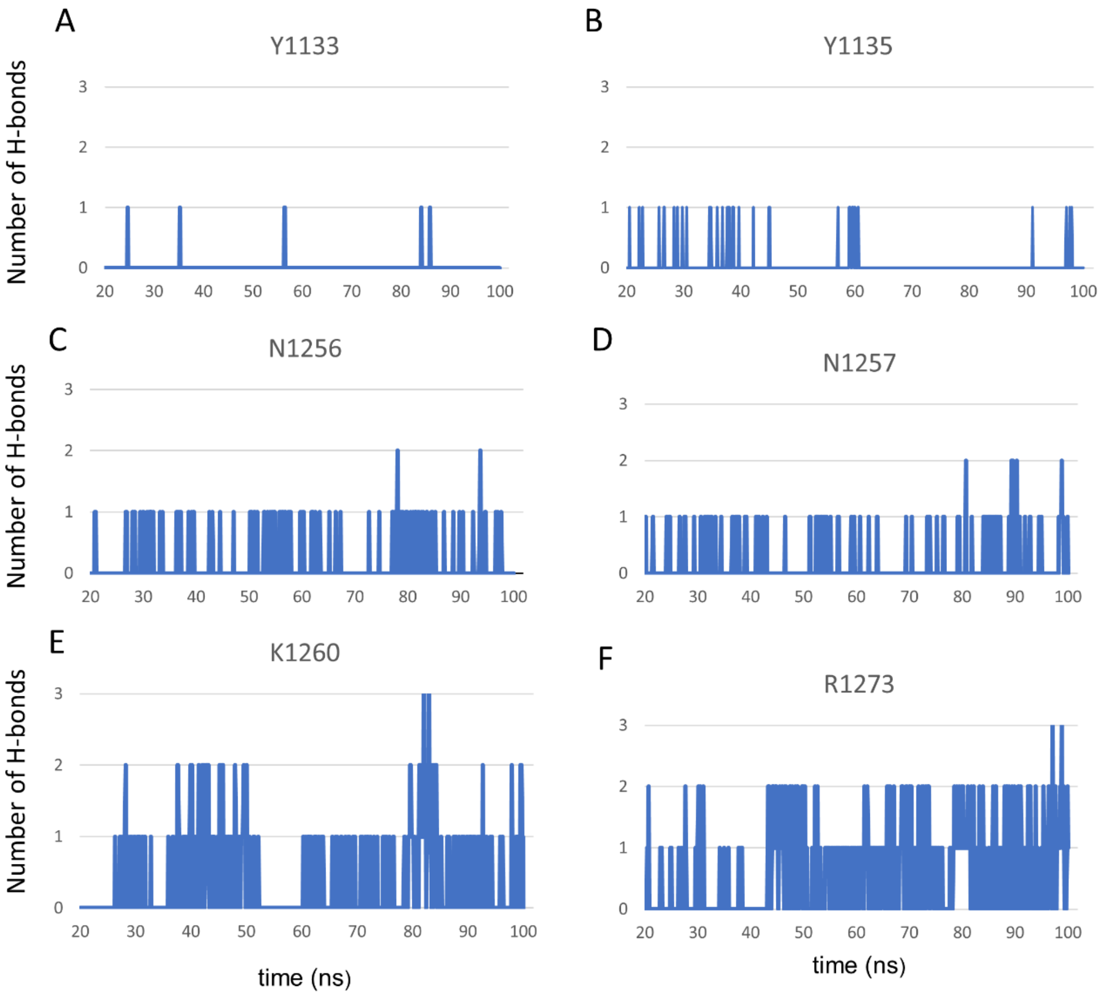


**Figure S6:** Plots showing the number of H-bonds formed between the sialic acids of gangliosides and Y1133 **(A)**, Y1135 **(B)**, N1256 **(C)**, N1257 **(D)**, K1260 **(E)** and R1273 **(F)**.

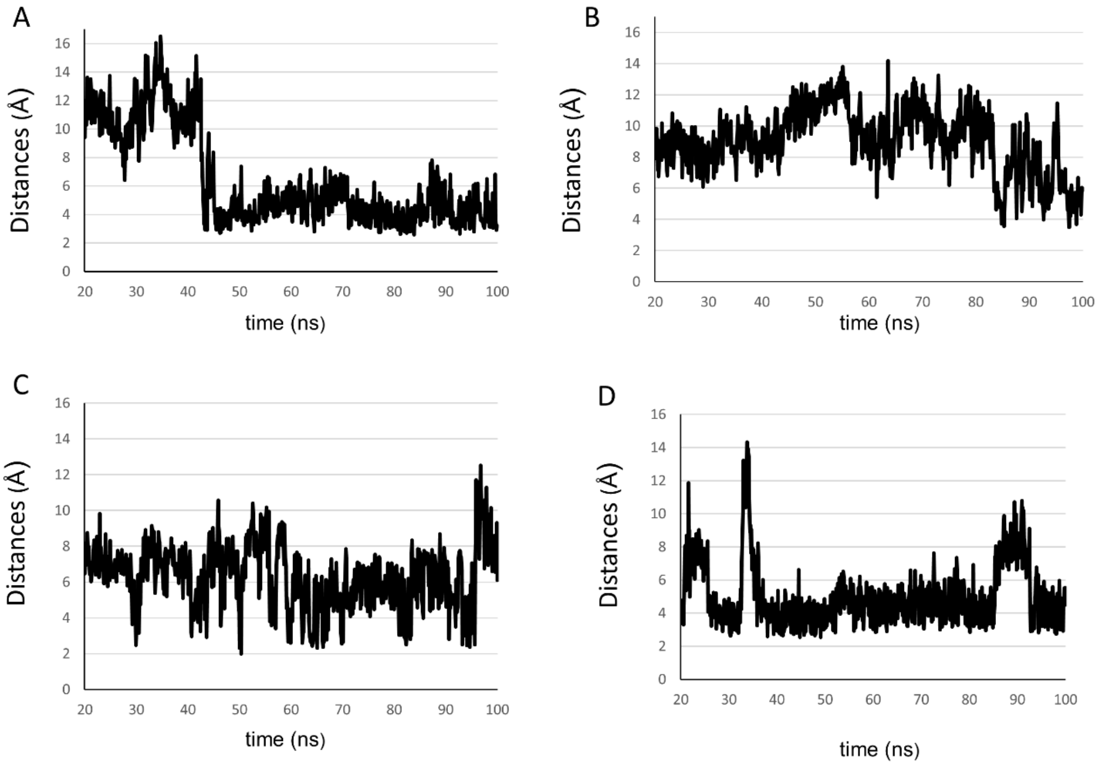


**Figure S7:** Plots showing the distances between the sialic acids of gangliosides and the aromatic ring of Y1133 **(A)**, Y1135 **(B)**, H1253 **(C)** and F1255 **(D)**.

| | Total energy of interaction (kJ/mol) at: | | | |
|---|---|---|---|---|
| | 40 ns | 60 ns | 80 ns | 100 ns |
| SV2C/1g | -19.19 | -33.77 | -16.79 | -3.87 |
| SV2C/2g | -121.69 | -118.44 | -115.58 | -104.40 |

**Table S1:** Total interaction energy of N559g in SV2C/1g system (first line) or SV2C/2g system (last line) at 40, 60, 80 and 100 ns.
